# Filamentous aggregates are fragmented by the proteasome holoenzyme

**DOI:** 10.1101/467068

**Authors:** Rachel Cliffe, Jason C Sang, Franziska Kundel, Daniel Finley, David Klenerman, Yu Ye

**Affiliations:** Department of Chemistry, University of Cambridge, Lensfield Road, Cambridge CB2 1EW; Department of Cell Biology, Harvard Medical School, Longwood Avenue, Boston 02115 MA

**Author notes:** Equal contribution.

## Abstract

Filamentous aggregates (fibrils) are regarded as the final stage in the assembly of amyloidogenic proteins and are formed in many neurodegenerative diseases. Accumulation of aggregates occurs as a result of an imbalance between their formation and removal. Although there have been numerous studies of the aggregation process *in vitro*, far fewer studies of aggregate disassembly and degradation are available. Here we use single-aggregate imaging to show that large fibrils assembled from full-length tau are substrates of the 26S proteasome holoenzyme, which fragments them into small aggregates. TEM further revealed that these small aggregate species had no distinct structure. The intact proteasome holoenzyme is required to effectively target fibrils. Interestingly, while degradation of monomeric tau was not inhibited by ATPγS, fibril fragmentation was predominantly dependent on the ATPase activity of the proteasome. The proteasome holoenzyme was also found to target fibrils assembled from α-synuclein (αS), suggesting that its fibril fragmenting function may be a general mechanism. The fragmented species produced by the proteasome showed significant toxicity to human cell lines compared to intact fibrils. Together, our results indicate that the proteasome holoenzyme possesses a novel fragmentation function that disassembles large fibrils into smaller and more cytotoxic species.

## Introduction

Protein aggregation is often associated with neurodegenerative disorders and aging-related dementia[1]. Alzheimer’s and Parkinson’s diseases are common dementia-like disorders involving the aggregation of distinct amyloidogenic proteins tau and α-synuclein (αS), respectively[2,3]. The protein tau is suggested to participate in the assembly and stability of microtubules but has also been associated with other functions[4]. αS is able to interact with phospholipids and vesicles and believed to be involved in cellular vesicle trafficking and neurotransmitter release[5]. Both tau and αS are largely intrinsically disordered when not associated with other proteins[6,7].

Amyloidogenic proteins have a propensity to misfold and oligomerize[8]. As oligomers grow in size by addition of protein monomers, conformational changes that are associated with increased stability take place, eventually resulting in a highly ordered and filamentous arrangement of aggregates that are no longer soluble in the physiological environment[9]. How these aggregates relate to toxicity leading to cell death and ultimately cause pathological disorders remain disputed, though distinct types of aggregates have been found to impede cellular signaling and compromise the integrity of neuronal functions[10-13]. The size and the type of aggregates may also determine how they are processed by cells and either targeted for degradation or sequestered at distinct cellular sites (e.g. [14]).

Although the formation of aggregates has been researched extensively, little is known about their removal. Aggregate degradation via both the proteasomal and the lysosomal systems has been described in the literature (e.g.[15,16]). Whereas larger aggregates are believed to be cleared by lysosomes, removal of smaller oligomers has been attributed to the proteasome[17]. The 26S proteasome holoenzyme is an abundant multisubunit protein complex responsible for the regulation of many key signaling pathways and general cell homeostasis[18]. This complex consists of a cylindrically shaped 20S core particle (CP) and one or two 19S regulatory particles (RP) that cap the CP at either end[19] (Modeled in **SFigure 1a**). Degradation activity is provided by several proteases within the interior of the CP, while the RP is responsible for the recognition of ubiquitin(Ub)-modified substrates, which are subsequently unfolded and translocated into the CP[20]. Six ATPases arranged in a hexameric ring are found within the base of the RP, which couples ATP hydrolysis to substrate unfolding and translocation through its channel pore. In cells, both tau and αS have been reported to be degraded by the proteasome[21,22], while aggregates assembled from these proteins were not found to be targeted by the proteasome *in vitro* (e.g. [23,24]). It is further possible that distinct aggregate conformations of sufficient size and stability may be recognized but not processed by the proteasome and thus inhibit its activity, as has been suggested for tau, αS, amyloid-β and prion aggregates[23,25-27].

Here we use single-aggregate total internal reflection fluorescence (TIRF) microscopy to show a novel aggregate fragmentation function of the proteasome holoenzyme, which targets fibrils in a Ub-independent manner. Fibrils assembled from full-length tau were predominantly removed by the proteasome in an ATP-dependent manner, while inhibiting the proteolytic activity of the proteasome had negligible effect on the fragmentation function. Fragmentation was further confirmed by transmission electron microscopy (TEM), revealing a species that resembled amorphous aggregates following proteasome treatment. This aggregate species was more toxic to cultured mammalian cells than fibrils, triggering a significant level of cell death. Our findings were further confirmed using αS fibrils, suggesting that this fragmenting function is not restricted to targeting tau fibrils. Together our findings demonstrate a novel ability of the proteasome holoenzyme to disassemble fibrils and its activity may be regulated through altering the physiological ratio of the holoenzyme to the free proteasomal core and regulatory particles.

## Results

### Proteasome holoenzyme degrades monomeric tau

To study how filamentous aggregates (fibrils) may be processed by mammalian proteasomes (**SFigure 1a**), we purified the holoenzyme or the RP separately from established HEK293T cells[28](see **Materials and Methods**). The purity and integrity of the proteasomes were confirmed using SDS-PAGE and TEM (**SFigure 1b** and **c**). Untagged recombinant full-length tau (isoform 0N4R, modeled in **SFigure 2a**) containing a single Pro274Ser substitution was purified to apparent homogeneity (**SFigure 2b**) and subjected to proteasomal degradation. The Pro274Ser substitution enhances tau aggregation and is commonly used in tauopathy models[29]. Monomeric tau was degraded by the holoenzyme, as demonstrated by the loss of substrate band intensity over time (**SFigure 2c**). Degradation by the holoenzyme was efficiently inhibited by 50 µM Velcade alone or an inhibitor cocktail (50 µM each of Velcade, MG132 and Carfilzomib; all of which target proteasomal proteases), but not by the replacement of ATP with a slowly hydrolysable ATP analogue, ATPγS (**SFigure 2c**). This result suggests that the degradation of monomeric tau is dependent on the proteolytic but not the ATPase-catalysed unfolding-translocation activity of the proteasome and that Velcade is sufficient to fully inhibit proteasomal degradation of tau proteins.

### Tau fibrils are fragmented in presence of the proteasome holoenzyme

We next tested whether aggregates assembled from tau may also be targeted by the proteasome holoenzyme. Fibrils assembled from tau could be reproducibly obtained at similar levels after 24 hrs of aggregation reaction following established protocols (e.g.[30]). Aggregated tau samples were treated with the proteasome or an ATP-containing buffer control and subsequently mixed with a second solution containing pentameric formylthiophene acetic acid (pFTAA, **Figure 1a**), a fluorophore that emits fluorescence upon binding to amyloid structures in aggregates[31]. We further established an approach to detect aggregated proteins directly on a glass coverslip surface (**Figure 1b**, *left*). Our approach does not require prior labeling of tau proteins and permits fluorophores in solution to reversibly bind the aggregates, thereby prolonging imaging lifetime. Aggregates were imaged on a custom-built fluorescence TIRF microscope (**SFigure 3a**) and analyzed using custom-written scripts that we have developed to assess individual aggregate size and fluorescence intensity, which in turn reflects the level of amyloid structures present (**SFigure 3b** and **Materials and Methods**).

**Figure 1.**
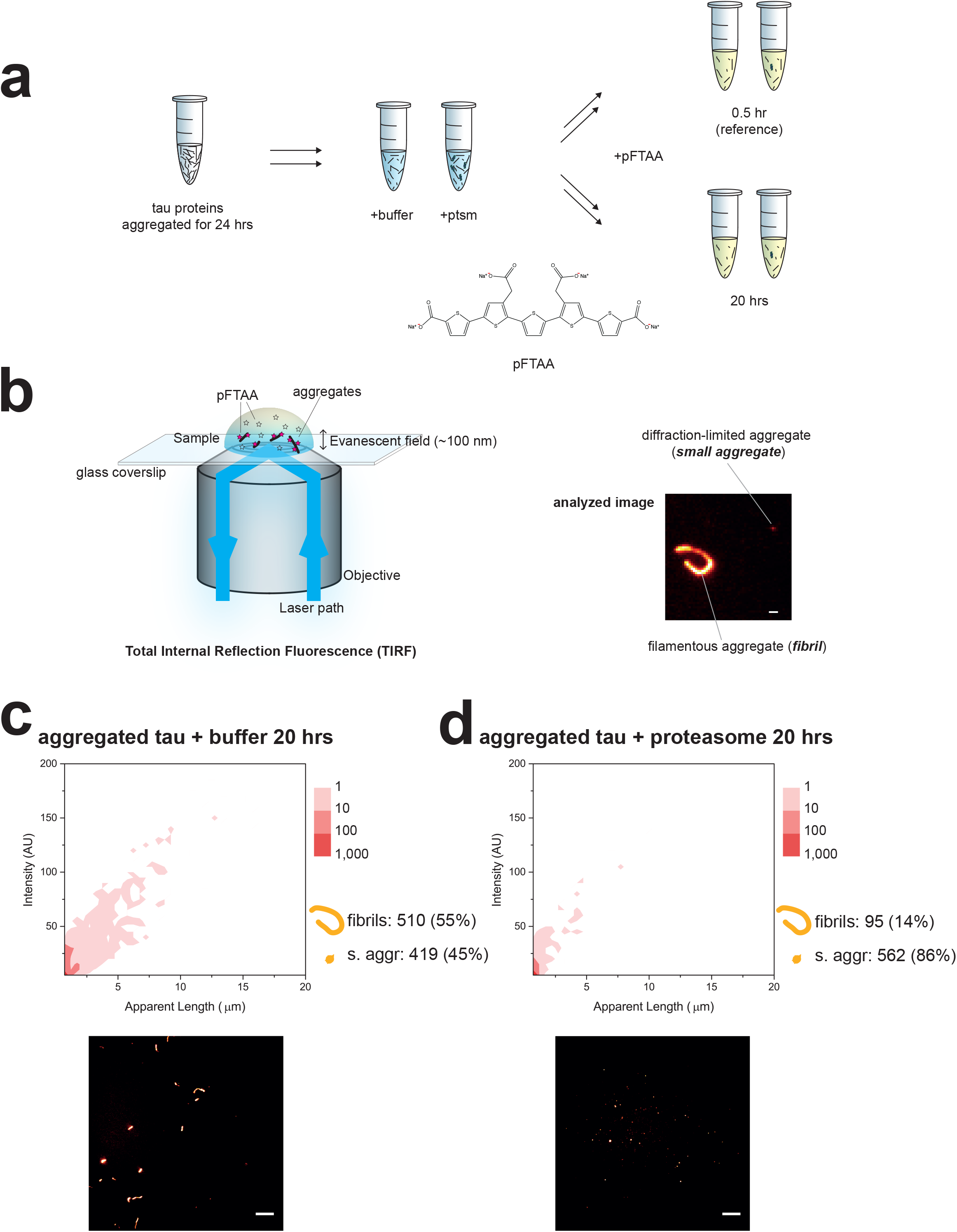
Imaging filamentous aggregates (fibrils) with fluorescence TIRF microscope. **(a)** Recombinant full-length tau was aggregated for 24 hrs and aliquots were taken and mixed with the proteasome in an ATP-containing Proteasome buffer or with the buffer only as a control. After 0.5 hr (starting reference) and 20 hrs of incubation, each reaction was diluted in an imaging buffer containing pFTAA. The chemical structure of pFTAA, which binds amyloid structures, is shown. **(b)** Samples were placed on a glass coverslip, excited with a 488 nm laser and imaged on a custom-built TIRF microscope (see also **SFigure 3**). A typical ‘fibril’ (length > 1 µm) and diffraction-limited ‘small aggregate’ (length < 1 µm) are shown. Scale bar represents 1 µm. **(c)** A large amount of fibrils remained present after incubation with the buffer alone, while **(d)** treatment with the proteasome holoenzyme resulted in loss of fibrils and an increase in small aggregate count (depicted next to the 2D plot). The length of aggregates is plotted against the fluorescence intensity of pFTAA, where the frequency is color-coded in the 2D plots. A processed image from each reaction is shown below respective plots. Scale bars represent 10 µm. Results of three biological repeats (n = 3) performed independently using different protein preparations of tau and proteasome were combined into each plot. The standard deviation between repeats was less than 20%.

The size (apparent length) of individual pFTAA-positive aggregates as detected by TIRF was plotted against their fluorescence intensity and presented in two-dimensional (2D) graphs. The level of amyloid structures defined in each aggregate increases proportionally with aggregate size (**SFigure 3c**). Based on the contours of aggregates, we will refer to the large aggregates (length >1 µm) with distinct shapes and high amyloid structure content as *fibrils* and those with indistinct morphology due to the diffraction limit (length < 1 µm) as *small aggregates* (**Figure 1b** *right* and **SFigure 3d**).

Large fibrils (up to 15 µm in length) assembled from tau proteins could still be detected even after 20 hrs of incubation in the degradation buffer without the proteasome. These fibrils constituted about 55% of all aggregates (510 fibril and 419 small aggregate counts, **Figure 1c**). In comparison, incubation with the proteasome holoenzyme quantitatively removed tau fibrils (95 counts, 7% of total), while the level of small aggregates increased (562 counts, **Figure 1d**). The standard deviation from the mean of experiments was less than 20% for both fibrils and small aggregates. Proteasomes alone did not bind pFTAA and therefore could not contribute to any fluorescence signals detected (**SFigure 4a**). An ATP regeneration system was added to both the control and proteasome-treated samples to maintain the ATP concentration during the assay (see **Materials and Methods**).

### Fibrils are targeted by ATP-dependent proteasomal activity

An equilibrium may potentially exist between fibrils, soluble oligomers and monomers, the last of which could be degraded by the proteasome and thus lead to an equilibrium shift favoring fibril disassembly. To validate that fibrils were being targeted by the proteasome, we repeated the experiments in **Figure 1c** and **d** after soluble tau monomers and oligomers were separated from aggregated proteins by centrifugation (see **Materials and Methods**). After removing the supernatant, the pellet containing the fibrils was resuspended with a fresh buffer. Following incubation in the buffer control for 20 hrs, fibrils up to 15 µm in length (407 counts, or 54% of total, **Figure 2a**) were still detected at a similar level as in **Figure 1c**. As expected, incubation with the holoenzyme led to fibril fragmentation, resulting in a drop in the level of fibrils (155 counts, 12% of total) and an increase in the number of small aggregates (891 counts, **Figure 2b**). These results indicate the presence of a novel proteasomal function that fragments tau fibrils. Plausibly, the proteasome may fragment a single fibril into many small aggregates, at least some of which may be detected by TIRF imaging. The increase in the level of small aggregates here is in agreement with **Figure 1** and suggests that fibrils may have been fragmented into smaller species of no more than 1 µm in size. We further addressed the possibility of contamination by canonical chaperones that may be responsible for fibril fragmentation; the fragmenting activity was unaffected by inhibitors of HSP70, HSP90 or VCP/p97 (**SFigure 4b-e**). Therefore, the loss of fibrils and the increase in small aggregates appear to be results of proteasomal action.

**Figure 2.**
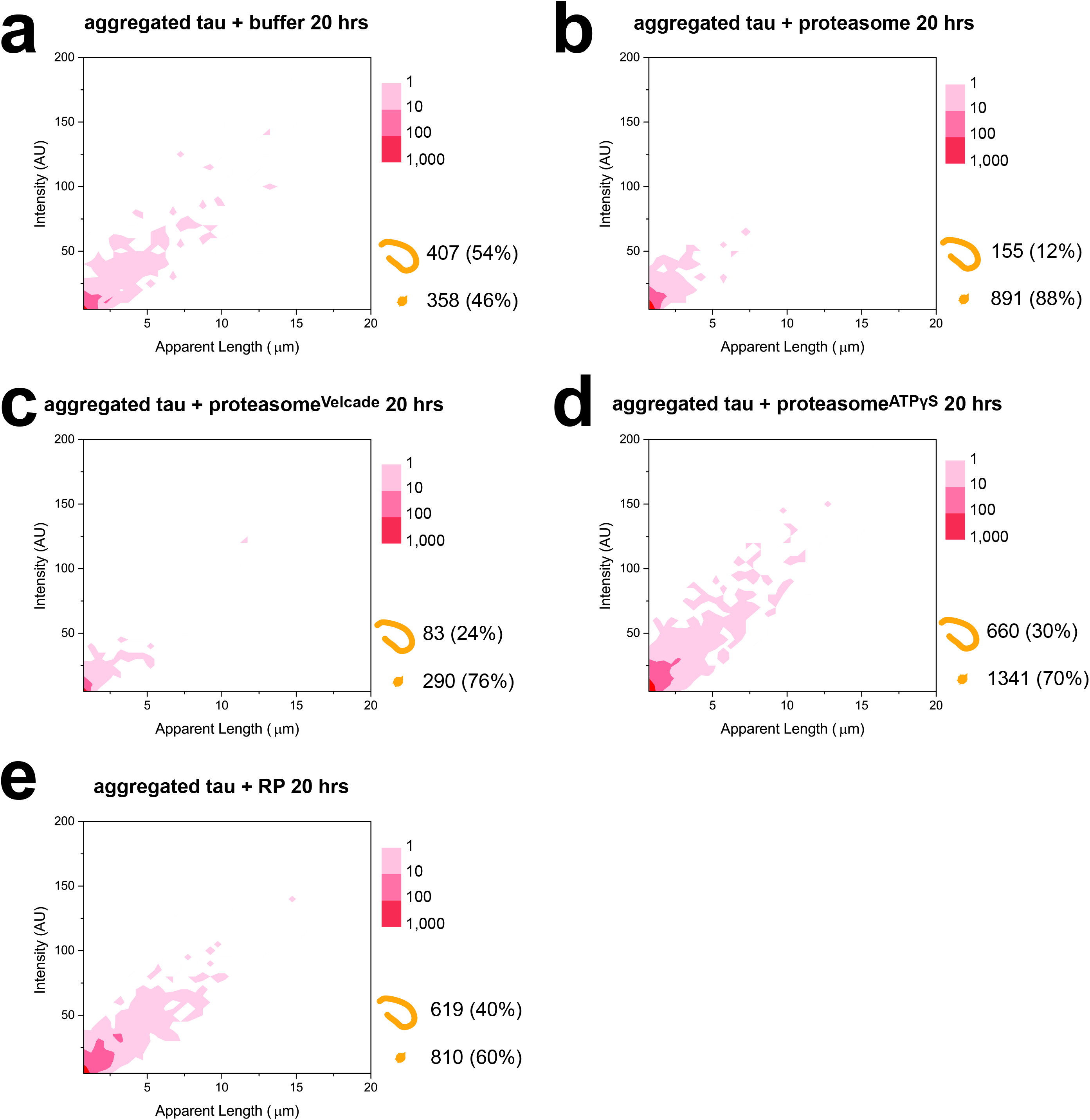
Fragmentation of fibrils in the absence of soluble tau proteins. Aggregated tau samples were centrifuged and the pellet resuspended in fresh Proteasome buffer followed by incubation with **(a)** buffer control or **(b)** the proteasome holoenzyme for 20 hrs and subsequently imaged and presented as described in **Figure 1**. Proteasome holoenzyme pre-treated with **(c)** Velcade (proteasome^Velcade^) or **(d)** ATPγS (proteasome^ATPγS^) were subsequently incubated with aggregated tau as in **b**. Instead of the holoenzyme, fibrils were also incubated with **(e)** regulatory particles (RP) and analyzed as before. Combined results of three independent experiments (n = 3) are shown here for all panels.

To gain insights into the proteasomal mechanisms responsible for this fibril-fragmenting function, we repeated the same assay using holoenzymes pre-treated with either Velcade or ATPγS. Fragmentation was largely observed (83 counts of remaining fibrils) even after inhibition of proteolytic activity (proteasome^Velcade^, **Figure 2d**), further excluding the scenario that proteolytic degradation of tau proteins mediates the loss of fibrils. The number of small aggregates (290 counts) remained at a similar level as control in **Figure 2a**. In contrast, compromising the ATP-dependent activity (proteasome^ATPγS^) impeded the fragmenting function, leaving the fibril level largely unchanged (660 counts, **Figure 2e**). The lower ratio of fibrils (30%) when incubated with proteasome^ATPγS^ is due to the apparent level of small aggregates detected (1341 counts). These observations together imply that the fragmentation of fibrils relies predominantly on the ATPase activity the proteasome.

### Integrity of proteasome holoenzyme required for fibril fragmentation

Since fibril fragmentation was mainly dependent on proteasomal ATPase activity, we attempted to independently verify our observations using purified free RPs. Unexpectedly, no significant fibril fragmentation was detected in the presence of RP (**Figure 2e**), and the level of fibrils (619 counts, or 40% of total) remained largely similar to **Figure 2a**. This result suggests that the integrity of the proteasome holoenzyme may be required to couple functions that are required for efficient fibril fragmentation.

### Distinct activities of proteasome holoenzyme prevent tau aggregation

While the fibril-fragmenting function of proteasome holoenzymes may be relevant to target aggregates that are already assembled, e.g. in a scenario when aggregates enter the host cell from the extracellular environment (e.g. [32-34]), the proteasomal mechanisms that are involved in clearing cytosolic misfolded proteins to prevent intracellular aggregate formation may be distinct from its fibril-fragmenting function. To further validate the observations in **Figure 2** and mimic a physiological scenario where proteasomes are already present during the aggregation process, we attempted to assemble tau aggregates in the presence the holoenzyme. Without the proteasome, fibrils emerged 0.5 hr after aggregation start (**Figure 3a**), and a significant number of fibrils (994 counts or 34% of total, **Figure 3b**) was observed after 24 hrs of reaction. In comparison, aggregation in the presence of holoenzymes restricted the ratio of fibrils (13%, 141 counts) assembled after 24 hrs (**Figure 3c**). Selectively inhibiting the proteolytic or ATPase activities of the proteasome partly restored the fibril level (704 and 978 counts, respectively, **Figure 3d** and **e**), suggesting that both activities are involved in impeding fibril formation from protein monomers. It therefore appears that the proteasomal mechanisms observed here, which are involved in reducing tau aggregation, are distinct from the fibril-fragmenting function in **Figure 2**, where the proteasomes were introduced to pre-assembled fibrils. Interestingly, proteasome^ATPγS^ incubation enabled both more and large fibrils (over 10 µm) to form compared to proteasome^Velcade^, consistent with a central role of the ATPase activity in fibril disassembly.

**Figure 3.**
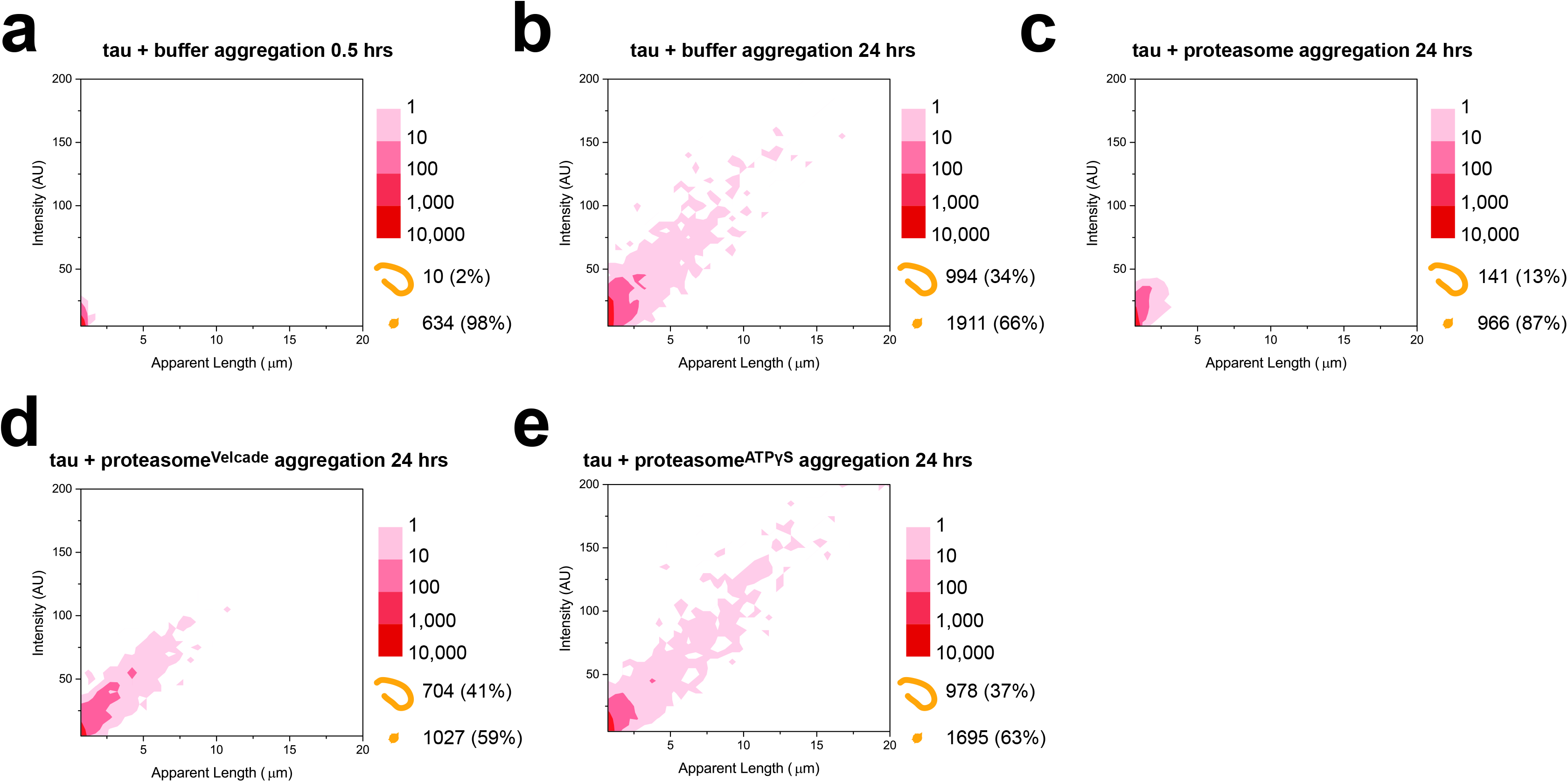
Aggregation of tau in the presence of proteasome holoenzymes. Monomeric tau proteins at 2 µM final concentration were mixed with the Proteasome buffer and imaged after **(a)** 0.5 hr or **(b)** 24 hrs, showing a substantial increase in both the fibril and small aggregate levels. Aggregation of tau in the presence of 40 nM final concentration of **(c)** untreated-, **(d)** Velcade- or **(e)** ATPγS-treated proteasome holoenzyme, measured after 24 hrs of incubation. Each plot contains the cumulative data from three independent measurements (n = 3).

### TEM detects amorphous structures following proteasome treatment

The fragmentation function of the proteasome in **Figure 2** was further independently validated using TEM. Fibrils in the absence of the proteasome remained intact after incubation with buffer control (**Figure 4a**). These fibrils were lost following proteasome treatment, and only unstructured clusters of proteins resembling amorphous aggregates were detected (**Figure 4b**), which were not present without proteasome treatment. Due to the intrinsic properties of uranyl acetate, which stains aggregates as well as proteasomes, we repeated these experiments using immunogold labeling with an anti-tau antibody to confirm the presence of tau proteins within these amorphous structures. This approach selectively labeled fibrils in the control sample (**Figure 4c**), as well as the fragmented species of amorphous structures after proteasome treatment (**Figure 4d**). These data indicate that tau-containing amorphous structures are formed following fibril fragmentation by the proteasome. Because the dimensions of these amorphous aggregates are mostly less than 1 µm, they are likely to have contributed to the small aggregates observed in **Figure 2**.

**Figure 4.**
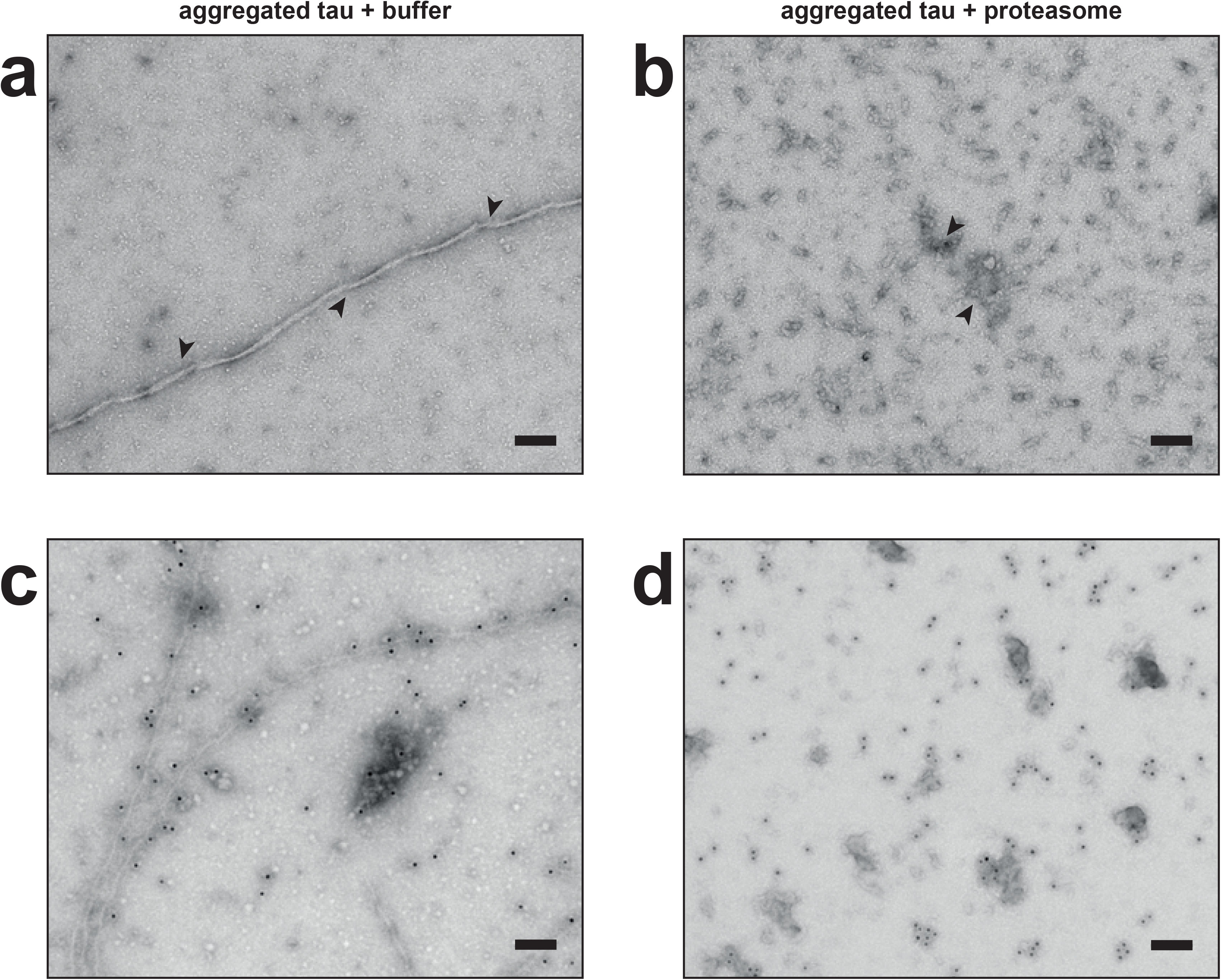
Disordered aggregates of amorphous structures detected by transmission electron microscope (TEM) after fragmentation by the proteasome. Aggregated tau samples prepared as in **Figure 2** were incubated with **(a)** an ATP-containing buffer or **(b)** proteasome holoenzyme for 20 hrs, stained with uranyl acetate and imaged by TEM. Arrows indicate typical aggregated structures. Aggregated samples incubated with **(c)** the buffer control or **(d)** the proteasome were immunolabeled with an anti-tau antibody, stained with uranyl acetate and imaged by TEM. Scale bars represent 100 nm. Representative images are shown of at least three independent repeats.

Intact proteasome holoenzymes could be detected under TEM after overnight incubation with the sample (see **Figure 4b**). This suggested that aggregated tau proteins did not affect the integrity of the holoenzyme. Consistently, we detected no quantitative inhibition of the proteasomal peptidase activity against a model substrate LLVY-AMC in the presence of fibrils or by a heparin-containing aggregation buffer (**SFigure 5a**), and incubating fibrils with the active proteasome holoenzyme for 20 hrs did not affect its ability to hydrolyze ATP (**SFigure 5b**). Although it may be possible that other proteasomal functions such as Ub-dependent degradation may be impaired as a result of prolonged interaction with aggregated tau proteins, any such inhibitory effect would not affect interpretation of our fragmentation data.

### Fibril fragmentation may increase cell lysis

Aggregates assembled from amyloidogenic proteins have been suggested to trigger cell stress and cytotoxicity, for example by disrupting lipid bilayers[35]. We hypothesized that the fibril-fragmenting function of the proteasome holoenzyme that resulted in a higher number of small aggregates would also lead to increased cytotoxicity, manifested through cell death (referred to as ‘cytotoxicity’ hereafter). To assess the cytotoxicity associated with fibril fragmentation, we incubated mammalian cells with untreated fibril samples or fragmented species and measured cell viability after 24 hrs. Viability was monitored using an established assay based on the release of cytosolic lactate dehydrogenase (LDH) into the extracellular media upon lysis. No lysis was detected after incubation with fibrils, indicating that viability of cells was not affected (**Figure 5a**, *column 1 from left*). In contrast, cells incubated with the fragmented species showed a substantially higher level of cell lysis (**Figure 5**, *column 2*). The reduced cell viability was not caused by the buffer or the proteasome alone, neither of which affected cell lysis (**Figure 5a**, *column 3 and 4*). In comparison, fibrils treated with free RP alone did not show a significant level of cell lysis either, in agreement with our *in vitro* data in **Figure 2e**.

**Figure 5.**
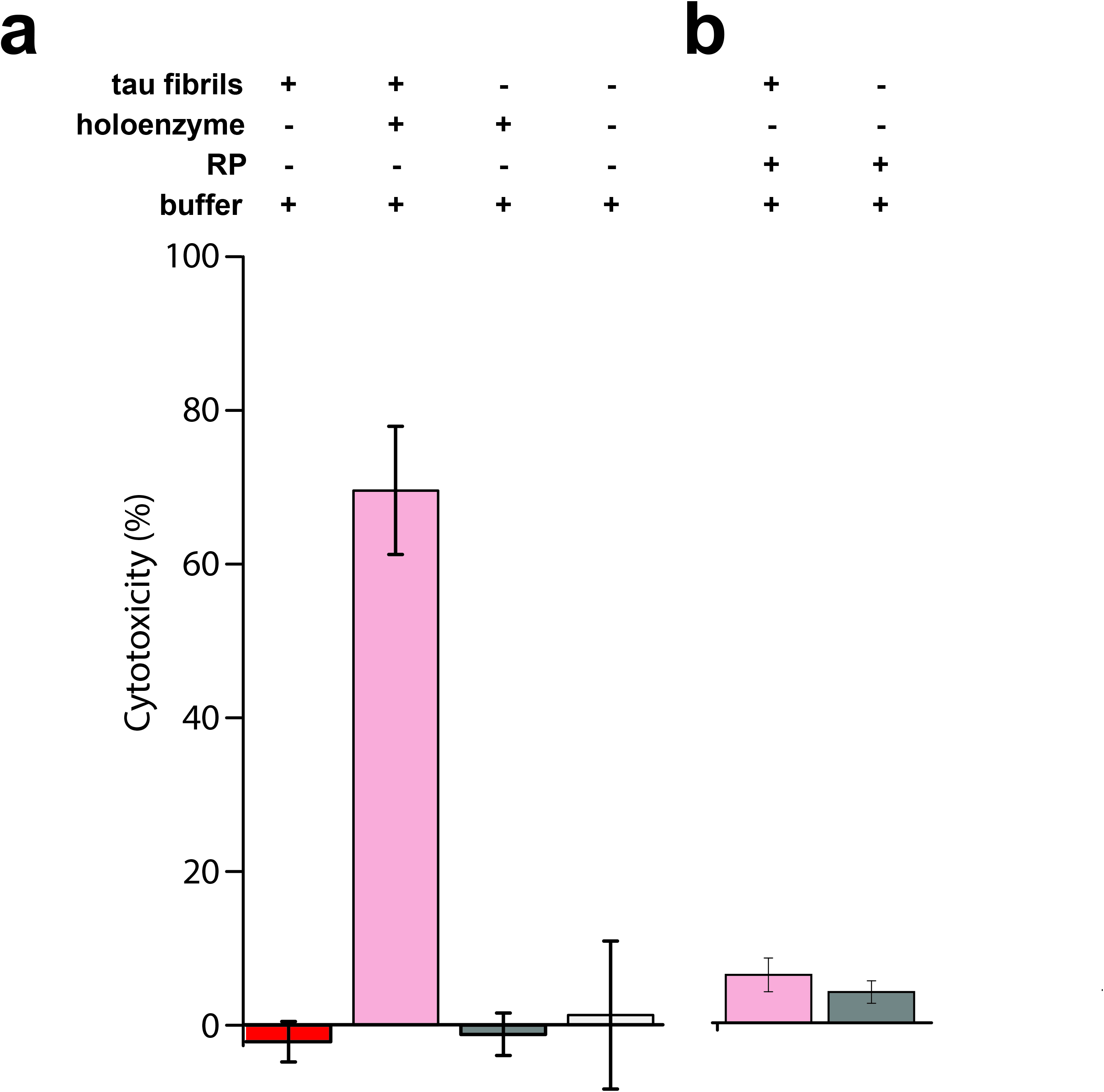
Fragmented species trigger cell lysis. **(a)** Lactate dehydrogenase (LDH) assay was used to detect lysis of HEK293 cells after incubation with either intact aggregated tau (*column 1*), the fragmented species (*column 2*), the buffer alone (*column 3*) or proteasome holoenzyme alone (*column 4*). **(b)** Cells were tested for level of lysis after incubation with fibrils treated with free RP (*column 5*) or with free RP alone as a control (*column 6*). Errors bars represent standard deviation of measurement from three independent experiments (n = 3).

Since the total number of aggregates increases after proteasome treatment, we addressed whether a higher concentration of fibrils would also lead to increased cell lysis. Cells incubated with fibrils alone at one-, ten- or hundred-fold higher concentration than in **Figure 5** did not cause any detectable cell lysis compared to the buffer or the tau monomer control (**SFigure 6a**). We further questioned whether any cytotoxic species might have escaped our detection during the centrifugation procedure, which separated insoluble fibrils from monomers and soluble aggregates. Neither the supernatant nor the pellet samples separated by centrifugation resulted in cell lysis (**SFigure 6b**). A positive control containing fibrils fragmented by sonication resulted in a high level of cell lysis (**SFigure 6c**), consistent with **Figure 5**. Together, these results indicate that the fibril fragmenting function of the proteasome may have a negative effect on cell viability.

### Conserved disassembly action of αS fibrils by the 26S proteasome

To test whether the fibril-fragmenting function may also be promiscuous and target other amyloidogenic proteins, we assembled fibrils from recombinant wild-type αS following previously established protocols (e.g.[36]). Purified monomeric αS is a substrate of the proteasome holoenzyme, and like tau its degradation is also dependent on the proteolytic but not the ATPase activity of the holoenzyme (**SFigure 7**). Fibrils assembled from αS could be reproducibly detected (1413 counts, or 28% of total) under the TIRF microscope after 20 hrs of incubation with buffer control (**Figure 6a**). These fibrils were fragmented by the proteasome holoenzyme, resulting in a decrease in the number of fibrils (1081 counts or 9%, **Figure 6b**). The lower ratio of fibrils is due to a substantial increase in the number of small aggregates (from 3682 to 10864 counts) following proteasome treatment. When untreated or proteasome-treated fibrils were stained with uranyl acetate and imaged by TEM, we detected smaller aggregate species of distinct structures following fragmentation (**Figure 6c** and **d**). These species were not amorphous and may account for the higher fluorescence signal detected by TIRF imaging after fragmentation of αS fibrils. Similarly to tau, fragmented αS entities were also significantly more toxic to cells (~28% cell lysis) than intact fibrils (~3% lysis, **Figure 6e**). Overall, the conservation of the fragmenting function against both tau and αS fibrils suggests that the proteasome holoenzyme may serve as a promiscuous fibril fragmentase and able to indiscriminately target fibril structures in general.

**Figure 6.**
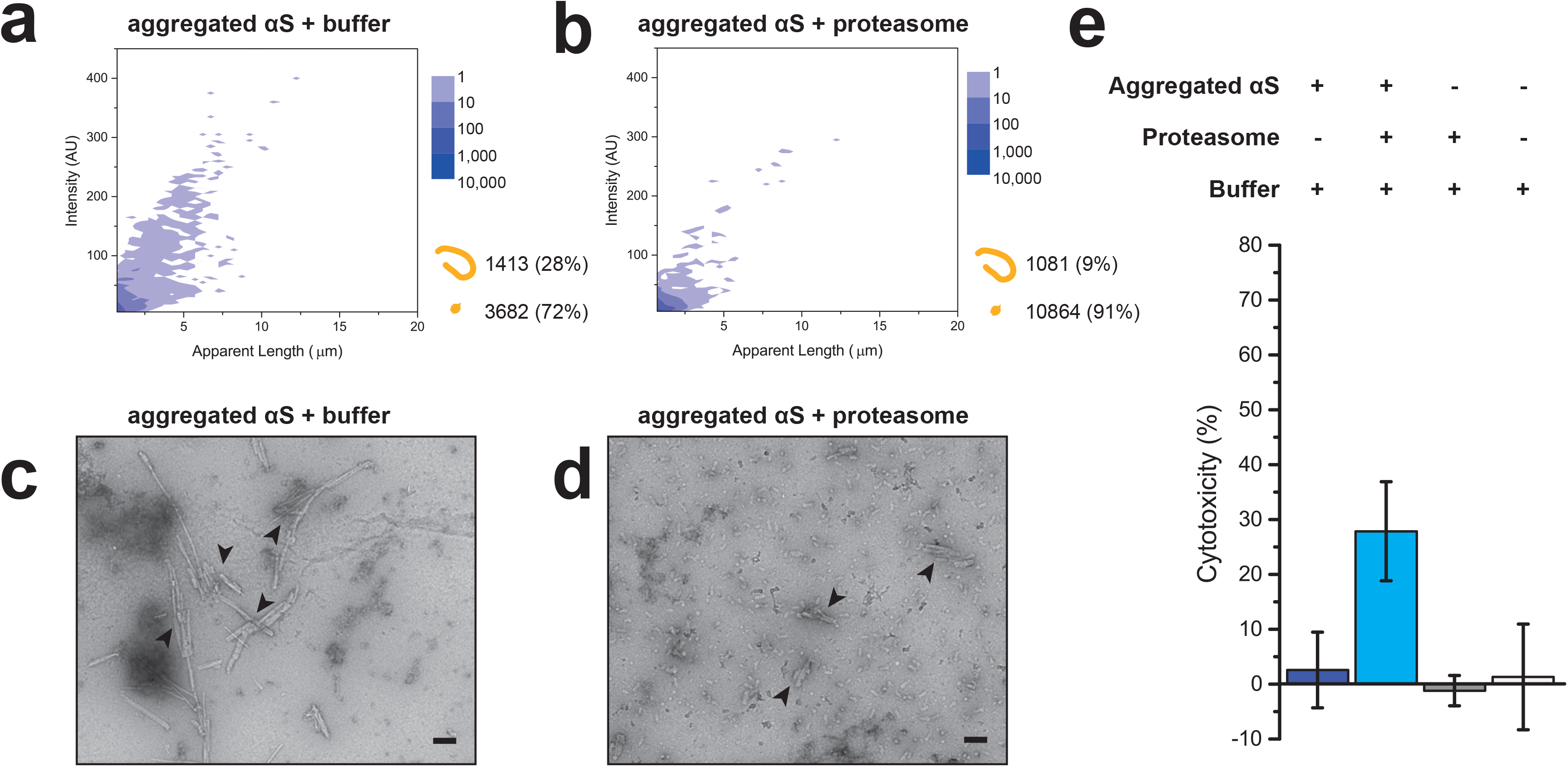
Proteasome fragments αS fibrils into cytotoxic species. Aggregates were assembled from αS monomers for 24 hrs and subsequently incubated with **(a)** an ATP-containing buffer or **(b)** proteasome holoenzyme for 20 hrs and presented in 2D plots as in **Figure 2**. Combined data of three independent repeats are shown (n = 3). **(c-d)** The assay was repeated with untreated or proteasome-treated aggregated αS for TEM imaging. Arrows indicate typical aggregated structures. Scale bars represent 100 nm. **(e)** HEK293 cells incubated with untreated - (*column 1*), proteasome-treated (*column 2*) samples of aggregated αS proteins, the buffer (*column 3*) or the proteasome alone (*column 4*) were tested for cytotoxicity using the LDH assay as in **Figure 5a**. All experiments have been independently repeated at least three times (n = 3) using fresh αS and proteasome preparations.

## Discussion

We have identified in this study a novel fibril-fragmenting function of the proteasome holoenzyme, which targets fibrils assembled from tau and αS proteins in an ATP-dependent manner and does not require its proteolytic activity. This novel fibril-targeting function makes the proteasome holoenzyme a structurally and functionally distinct fibril-fragmenting enzyme compared to other aggregate-targeting enzymes in the existing family of disaggregases such as Hsp104, which reverses aggregation of prion proteins[37]; and several HSP70-containing disaggregase complexes[38,39], one of which reported to target αS aggregates[39]. We further showed that this fragmenting function of the proteasome on pre-assembled fibrils is distinct to the proteasomal actions involved in preventing aggregation of misfolded proteins, which requires both the ATPase and proteolytic activities.

The proteasome has canonically been described to target oligomers and smaller aggregates in solution, whilst degradation of larger filamentous aggregates is performed by the lysosomal system[17]. We provide evidence in this study that amend this view, showing a novel ability of proteasome holoenzymes with molecular dimensions ~15 × 40 nm (pdb-id: 5GJR) to fragment fibrils up to ~20 µm in length independently of Ub modification. The recent molecular structure of tau filaments derived from an Alzheimer’s patient reveals extensive β-sheet stacking of tau monomers, suggesting a high degree of stability[40]. Our results therefore imply a remarkable ability of the proteasome to target and fragment stable structures several orders of magnitude larger.

Early *in vitro* studies have suggested an ATP-dependent chaperone-like function of the RP with affinity to denatured protein structures without Ub modification[41]. Recent *in vivo* studies have further found proteasomal recruitment to aggresomes in HEK cells[42,43] and to poly-GA aggregates in neurons[44], hinting a plausible proteasome involvement in targeting aggregates. Our findings in this study provide additional evidence suggesting that the ATP-dependent fibril-fragmenting function may also be driven by the unfolding or chaperone-like mechanism, perhaps when substrates cannot be a proteolytic target susceptible to direct degradation.

The current study has focused on the fibril-fragmenting function of proteasome holoenzymes on unmodified recombinant tau and αS. Under physiological settings, extensive posttranslational modifications have been identified on tau that affect its aggregation properties or are associated with pathological consequences[45,46]. Similar effects of various posttranslational modifications have also been reported for αS[47-49]. The range of aggregate species found *in vivo* that may be targeted by the fibril-fragmenting function of the proteasome, or alternatively those that are resistant to or inhibitory of this function, remain to be studied. Identification of the proteasome holoenzyme with a novel fibril fragmenting activity, in addition to its canonical role as a mediator of the degradation of misfolded proteins, could be an interesting amendment to the repertoire of cellular instruments targeting protein aggregation.

## Acknowledgement

The authors thank members of the Klenerman and Finley labs for reagents and useful discussions. YY is funded by a Sir Henry Wellcome Research Fellowship. Works in DK and DF labs are funded by grants from the NIH (GM043601) and from the EPSRC, respectively.

## Author contributions

YY and RC performed the experiments and analyzed the data. JCS wrote the script for image processing. FK provided protein reagents and assisted with experiments. YY, DK, DF designed the experiments and directed the research. YY conceptualized the project and prepared the manuscript.

## Materials and Methods

### Protein purification

Plasmids expressing untagged full-length tau (isoform 0N4R) containing a single Pro274Ser substitution or wild-type α-synuclein (αS) in pT7-7 vectors were transformed into BL21 cells. Protein expression was induced at an O.D. of 1.0 using 1 mM IPTG for 4 hrs at 20 °C. Cells were collected by centrifugation at 4000 × g and then resuspended in ice-cold Lysis buffer (50 mM MES pH 6.5, 2.5 mM TCEP, 1 mM AEBSF). Lysis of cells was carried out by sonication and the lysate was cleared by centrifugation at 23000 × g for 30 min at 4 °C. The pH of the cleared lysate containing tau was gradually reduced to 4.5 and incubated on ice for 10 min before repeating the centrifugation at 23000 × g to clear the lysate of precipitants. The supernatant containing tau was filtered and subsequently loaded onto a ResourceS ion exchange column (GE Healthcare) and eluted with a linear NaCl gradient. Eluted fractions containing tau were identified by SDS-PAGE and loaded onto Superdex 16/60 (GE Healthcare) gel filtration column for a final purification step (**SFigure 2b**). For αS purification, the cleared lysate was incubated in boiling water for 15 min followed by another centrifugation procedure at 23000 × g for 30 min at 4 °C. The αS supernatant was then loaded onto a ResourceQ anion exchange column and eluted with a linear NaCl gradient. Fractions containing αS were loaded onto Superdex 16/60. The eluted fractions from gel filtration were examined by SDS-PAGE and fractions judged pure were concentrated and flash-frozen.

Purification of mammalian proteasome holoenzymes or the free regulatory particle (RP) was carried out as described elsewhere[28]. Briefly, mammalian HEK293T cells stably expressing a fusion construct coding for Rpn11-His_6_-TEV-biotin were grown to confluency and collected using a cell scraper. The cells were resuspended in Proteasome buffer (50mM Tris [pH7.5], 5 mM ATP, 5 mM MgCl_2_)and lysed using a dounce homogenizer. Cell lysate was cleared by centrifugation at 1500 × g for 10 min and the supernatant was incubated with 2 ml NeutrAvidin beads overnight at 4 °C. On the next day, beads were washed with TB buffer (50mM Tris [pH7.5], 5 mM ATP, 5 mM MgCl_2_, 10% glycerol) and bound holoenzymes were eluted after 3 hrs of incubation with 6 µl TEV protease (Sigma) at 30 °C. For RP purification, the beads were first washed with TBN buffer (TB buffer containing 800 mM NaCl) to release the CP from the bound RP and then with additional TB buffer prior to the TEV protease cleavage step. Subsequent purification steps were the same as for the holoenzyme.

### Aggregation assays

Aggregation reactions for tau were set up at 2 µM final concentration in SSPE buffer (10 mM Na_3_PO_4_ (pH 7.4), 150 mM NaCl, 1 mM EDTA, 0.02% Sodium Azide) in the presence of 2 µM heparin (5000 Da, Fisher Scientific) at 37 °C[30]. For αS aggregation, protein was diluted to 70 µM final concentration in PBS buffer (Fisher Scientific) containing 0.02% Sodium Azide and performed under shaking conditions at 37 °C[36]. Both tau and αS were aggregated for 24 hrs and used immediately for subsequent reactions. All buffers used in our assays were pre-filtered with 0.02 µm filters.

Aggregation reactions in **Figure 3** contained 2 µM and 40 nM final concentration of tau and proteasome holoenzyme, respectively, in the Proteasome buffer with an ATP regeneration system (20 mM Creatine phosphate and 5 µM Creatine kinase final concentration) and initiated with 2 µM heparin at 37 °C as described above.

### Proteasome assays

Aliquots were removed from aggregation reactions after 24 hrs and incubated with the proteasome. The final reactions contained 8 µl of 200 nM proteasome, 8 µl of the aggregated tau or αS substrate, 5 µl of 10 × Proteasome buffer, 2.5 µl of an 20 × ATP regeneration system (2 M Creatine phosphate and 100 µM Creatine kinase) and ddH_2_O to make up 50 µl final reaction volume. The reactions were performed at 25 °C to avoid further aggregation. After 0.5 hr (used as a reference) and 20 hrs of incubation, 1 µl aliquot was removed from each reaction and serially diluted 50-fold in PBS buffer containing 30 nM pFTAA for TIRF imaging (see below). For experiments in **Figures 2 - 6**, prior to the proteasome treatment step, aggregated samples were centrifuged on a benchtop centrifuge at maximum velocity for 30 min. The supernatant was subsequently removed and resuspended with an equal volume of Proteasome buffer before incubation with the proteasome. In control reactions, the proteasome was replaced with an equal volume of the Proteasome buffer. The distinct catalytic activities of the proteasome were inhibited by pre-incubation with Velcade or ATPγS for 5 min in room temperature before mixing with the substrate. The final concentrations of Velcade and ATPγS used were 50 µM and 9 mM, respectively. No ATP was added to the buffer in reactions containing ATPγS. Proteasomes pre-treated with VER155008, Geldanamycin and NMS-873 were prepared the same way as Velcade and used at 50 µM final concentration.

### LLVY-AMC and Malachite Green assays

LLVY-AMC (Enzo Lifesciences) was used at 1 µM final concentration and incubated either with 40 nM final concentration of proteasome alone, with the proteasome in presence of 320 nM heparin or with 320 nM aggregated tau proteins. The sample was excited at 340 nm and emission detected over time on a fluorimeter (Cary Eclipse) at 440 nm.

ATPase kit containing the Malachite Green assay was purchased from Abcam (ab65622). For experiments in **Figure 5b**, 40 μl Proteasome assays set up as described above were mixed with 6 μl of the malachite green reagent and incubated for 15 min before measuring the O.D. at 650 nm on a microplate reader. Free phosphates were used to establish a linear standard curve to calculate phosphate concentrations from colorimetric readings.

### Degradation assays of protein monomers

Degradation assays of protein monomers were set up mixing 8 µl tau or αS at 2 µM or 70 µM, respectively, with 8 µl of 200 nM proteasome and H_2_O to make a final 50 µl reaction volume. At indicated timepoints, 6 µl of samples were removed and quenched with 6 µl LDS buffer and stored by flash-freezing for subsequent protein gel electrophoresis.

Protein samples were separated by 4-12% Bolt SDS-PAGE gels (Invitrogen) and transferred to PVDF membranes using Trans-blot Turbo (Biorad) semi-dry transfer system as per manufacturer’s protocol. Membranes were incubated with primary antibodies against tau (1E1/A6, Millipore) or αS (MJFR1, Abcam) overnight. Protein bands were detected using secondary anti-mouse or anti-rabbit antibodies tagged with Alexa647 (Invitrogen) and scanned on a Typhoon Imager.

### TIRF imaging

Prior to imaging, glass coverslips (0.13 mm thickness, VWR International) were cleaned with an argon plasma (PDC-002, Harrick Plasma) for 1 hr. A multi-well chambered coverslip (Sigma-Aldrich, GBL103350-20EA) was adhered to each glass coverslip to allow for imaging of multiple samples. For the imaging of αS samples, each well was coated with 0.01% poly-L-lysine (MW 150,000-300,000, Sigma-Aldrich) for 15 minutes, then washed three times with sterile-filtered PBS buffer. For tau samples, the glass coverslips were untreated. The imaging concentrations for tau and αS were 20 nM and 100 nM, respectively, of the calculated monomer concentration. Samples were imaged in the presence of 30 nM pFTAA dye (kind gift from Michele Goedert) in PBS buffer, using a home-built total internal reflection fluorescence (TIRF) microscope as shown in **SFigure 3a**. Images were recorded for 100 frames with 50 ms exposure time. For each sample, 9 fields of view were typically acquired and used for statistical analysis.

### Image analysis

Individual image data were averaged over all the frames by the average intensity projection at z-axis using ImageJ (National Institute of Health, USA) and then subjected to image processing. A custom-written MATLAB script (MathWorks) was used to analyze the averaged images. Individual images were first top-hat and bpass filtered to remove the camera noise and partitioned into a foreground and a background. To identify particles, the foreground was blurred using a 2D Gaussian filter with a threshold applied based on the original pixel intensity with a criterion of 2% intensity above the background, and then established boundaries for individual particles. The particle length was measured by thinning boundaries of individual particles and thus calculated with an image pixel size of 235 nm for our TIRF setup. To eliminate the background effect in intensity calculation, signal-to-background ratio (SBR) was introduced to correct pixel intensity, where each pixel’s SBR is defined as:

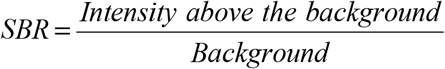

For a given particle, its corrected intensity is the sum of each pixel’s SBR values within its boundary.

### TEM imaging

Samples for TEM imaging were prepared as for TIRF imaging and applied onto a carbon-coated 400 mesh copper grid (Agar Scientific). Proteasome samples were applied at 100 nM final concentration. Samples were stained with 2% (w/v) uranyl acetate for 1 min and subsequently washed twice with ddH_2_O. TEM images were acquired using Tecnai G2 microscope (13218, EDAX, AMETEK) operating at an excitation voltage of 200 kV.

### Cytotoxicity assays

Fibrils treated with either the Proteasome buffer or with the proteasome holoenzyme (described in the Proteasome Assays section) were added in triplicate to confluent HEK293A cells grown in 96-well plates. The cells were subsequently incubated at 37 °C for 24 hrs in 200 µl final media volume, from which 100 µl was removed for LDH assay (Thermo Fisher). We followed manufacturer’s protocols and added 100 µl of the supplied Reaction Mixture, which contains the substrates for LDH activity detection, to the media. After 30 min, the reactions were quenched with the Stop Buffer (supplied with the kit) and the absorbance at 480 nm was measured on a plate reader. Lysis media supplied by the manufacturer was then added to each well to establish the maximum level of cell lysis. Assays were repeated at least three times using a new protein preparation each time. Both Velcade and ATPγS alone were found to be toxic to cells when added to the cells at the concentration used for proteasome inhibition, as significant levels of cell lysis was observed.

## Supplementary Figures

**SFigure 1.** Purification of mammalian proteasome assemblies. **(a)** A molecular model of the 26S proteasome holoenzyme (pdb-id: 5GJR) comprising one 20S core particle (CP, grey) capped at both ends with a 19S regulatory particle (RP, cyan). The ATPase subunits (light cyan) of the RP and the proteases of the CP (light grey) are highlighted. Rpn11 (magenta), a subunit of the RP, is modified at the C-terminus with a biotin-tag and used for affinity purification. **(b)** Purified RP (*left*) and proteasome holoenzyme (*right*) were resolved by SDS-PAGE and visualized by Coomassie staining. **(c)** TEM visualization of the holoenzyme (*left*) and the RP (*right*). Scale bars indicate 100 nm.

**SFigure 2.** Recombinant full-length tau monomers are degraded by the proteasome. **(a)** Model of the primary structure of full-length tau (isoform 0N4R, 383 Aas). The protein consists of an N-terminal protrusion region that does not participate in the assembled fibril structure, with distinct sequences that are acidic, basic and proline-rich. The tetra-repeat sequence within the C-terminal inclusion region is important to the assembly of tau aggregates. The tau proteins used in this study contain a single Pro274Ser substitution that enhances aggregation. **(b)** Pure recombinant tau proteins were obtained after a final gel filtration step. Fractions pooled for subsequent assays are marked. **(c)** *From left:* monomeric tau was incubated with the Proteasome buffer, the holoenzyme or holoenzyme pre-incubated with 50 µM Velcade, 30 mM ATPγS or an inhibitor cocktail. Aliquots from each reaction were quenched after 0, 5, 10 or 20 hrs, separated by SDS-PAGE and detected by immunoblotting against tau (clone 1E1/A6). The inhibitor cocktail contained 50 µM each of Velcade, MG132 and Carfilzomib.

**SFigure 3.** Imaging aggregated tau proteins using a custom-built TIRF microscope. **(a)** A model of the TIRF microscope set-up with a 488 nm laser (Cobolt MLD) directed to a 1.49 numerical aperture objective (APON60XO TIRF, Olympus) mounted on an inverted Nikon Eclipse Ti microscope. Fluorescence was collected by the same objective and separated from the returning TIRF beam by an appropriate dichroic (Semrock), and passed through appropriate emission filters (FF03-525/50-25, Semrock). Hardware was controlled using custom-written scripts for MicroManager (NIH) and images were recorded on an EM-CCD camera (Evolve 512Delta, Photometrics) with 235 nm image pixel size. **(b)** A schematic representation of the analysis workflow, where raw images were averaged over 100 frames, the fluorescence signals filtered and individual particles subsequently identified. Each particle is quantified by its size (apparent length as detected by TIRF) and fluorescence intensity (pFTAA binding). We define large filamentous aggregates (length > 1 µm) as ‘*fibrils*’ and near diffraction-limit aggregates (length < 1 µm) as ‘*small aggregates*’. Zoom-in shows a typical fibril next to a small aggregate according to the assigned criteria. Sizes of the scale bars are indicated below each row of images. **(c)** Each identified particle is plotted with respect to its length and fluorescence intensity. **(d)** The frequency of particles are binned together (length bin size 1, intensity bin size 5) and color-coded in the 2D plot. Regions are boxed according to where fibrils (cyan) and small aggregates (green) are found.

**SFigure 4.** Fragmentation of fibrils is specifically due to the proteasome. **(a)** The fluorescence level of proteasome holoenzyme alone in the absence of any aggregated sample is very close to the background level (*left*), with an insignificant amount of particles detected (*right*). (**b-e**) The proteasome holoenzyme was pre-treated with **(b)** an ATP-containing buffer control, **(c)** 50 µM Geldanamycin (HSP90 inhibitor), **(d)** 50 µM VER-155008 (HSP70 and HSC70 inhibitor) or **(e)** 50 µM NMS-873 (VCP/p97 inhibitor) and subsequently incubated with aggregated tau samples as in **Figure 2**. Cumulative data of three independent repeats (n = 3) are presented in each plot.

**SFigure 5.** No quantitative inhibition of the proteolytic and ATPase activities of the proteasome. **(a)** Proteasome holoenzymes were pre-treated with an ATP-containing buffer control (*green*), with the buffer containing 10 µM Heparin (*blue*), or with aggregated tau samples (*red*) in the same buffer. Proteolytic activity against a model fluorescent substrate LLVY-AMC was subsequently measured from its fluorescence emission. As a control, holoenzyme was pre-treated with 50 µM Velcade (grey). Error bars indicate the standard deviation from the mean value of three independent experiments. **(b)** ATP hydrolysis activity was measured using the malachite assay, which detects the concentration of free phosphates in the buffer. The level of free phosphates detected in the buffer alone (light grey), holoenzyme alone (brown) or holoenzyme with aggregated tau (magenta) incubated for 20 hrs as in **Figure 2** are shown.

**SFigure 6.** Aggregation buffers and tau fibrils are not toxic to cells. **(a)** Monomers (*column 1*) and fibrils (*column 2-4*) were incubated with the cells at ~50 nM as in **Figure 5**, at ten- and hundred-fold of the incubation concentration (10 × and 100 ×). **(b)** Cells incubated with SSPE buffer (*column 5*) or SSPE buffer containing heparin (*column 6*) used for tau aggregation were incubated and tested for cell lysis. The pellet (containing tau fibrils, *column 7*) or the supernatant fraction (containing soluble aggregates and monomers, *column 8*) were added to cells and tested for cell lysis. **(c)** Fibrils prepared as in **Figure 5** were sonicated briefly and centrifuged to separate insoluble fibrils before the supernatant was added to cells at the same calculated concentration as in **Figure 5**.

**SFigure 7.** Recombinant full-length αS monomers are degraded by the proteasome. **(a)** Monomeric αS incubated with buffer control (*left*) or the holoenzyme (*right*). **(b)** *From left:* Velcade-, ATPγS, or inhibitor cocktail-treated holoenzymes were incubated with monomeric αS. Aliquots from each reaction were taken at 0, 3, 5, 10 or 20 hrs after incubation, quenched and separated by SDS-PAGE. Changes in the αS level were detected by immunoblotting against αS (clone MJFR1). Experiments performed as in **SFigure 2c**. Representative data of at least three independent repeats.

## References

1. Goedert M (2015) Alzheimer’s and Parkinson’s diseases: The prion concept in relation to assembled Aβ, tau, and α-synuclein. Science 349: 1255555.

2. Iqbal K, Alonso ADC, Chen S, Chohan MO, El-Akkad E, Gong C-X, Khatoon S, Li B, Liu F, Rahman A, et al. (2005) Tau pathology in Alzheimer disease and other tauopathies. Biochim Biophys Acta 1739: 198–210.

3. Lashuel HA, Overk CR, Oueslati A, Masliah E (2013) The many faces of α-synuclein: from structure and toxicity to therapeutic target. Nat Rev Neurosci 14: 38–48.

4. Wang Y, Mandelkow E (2016) Tau in physiology and pathology. Nat Rev Neurosci 17: 5–21.

5. Bendor JT, Logan TP, Edwards RH (2013) The Function of α-Synuclein. Neuron 79: 1044–1066.

6. Theillet F-X, Binolfi A, Bekei B, Martorana A, Rose HM, Stuiver M, Verzini S, Lorenz D, van Rossum M, Goldfarb D, et al. (2016) Structural disorder of monomeric α-synuclein persists in mammalian cells. Nature 530: 45–50.

7. Schwalbe M, Ozenne V, Bibow S, Jaremko M, Jaremko L, Gajda M, Jensen MR, Biernat J, Becker S, Mandelkow E, et al. (2014) Predictive atomic resolution descriptions of intrinsically disordered hTau40 and α-synuclein in solution from NMR and small angle scattering. Structure 22: 238–249.

8. Soto C (2003) Unfolding the role of protein misfolding in neurodegenerative diseases. Nat Rev Neurosci 4: 49–60.

9. Spillantini MG, Goedert M (2013) Tau pathology and neurodegeneration. Lancet Neurol 12: 609–622.

10. Goedert M, Spillantini MG (2017) Propagation of Tau aggregates. Mol Brain 10: 18.

11. Labbadia J, Morimoto RI (2015) The biology of proteostasis in aging and disease. Annu Rev Biochem 84: 435–464.

12. Haass C, Selkoe DJ (2007) Soluble protein oligomers in neurodegeneration: lessons from the Alzheimer’s amyloid beta-peptide. Nat Rev Mol Cell Biol 8: 101–112.

13. Selkoe DJ (2004) Cell biology of protein misfolding: the examples of Alzheimer’s and Parkinson’s diseases. Nat Cell Biol 6: 1054–1061.

14. Woerner AC, Frottin F, Hornburg D, Feng LR, Meissner F, Patra M, Tatzelt J, Mann M, Winklhofer KF, Hartl FU, et al. (2015) Cytoplasmic protein aggregates interfere with nucleo-cytoplasmic transport of protein and RNA. Science 351: 173–176.

15. Wang Y, Mandelkow E (2012) Degradation of tau protein by autophagy and proteasomal pathways. Biochem Soc Trans 40: 644–652.

16. Webb JL, Ravikumar B, Atkins J, Skepper JN, Rubinsztein DC (2003) Alpha-Synuclein is degraded by both autophagy and the proteasome. Journal of Biological Chemistry 278: 25009–25013.

17. Rubinsztein DC (2006) The roles of intracellular protein-degradation pathways in neurodegeneration. Nature 443: 780–786.

18. Schmidt M, Finley D (2013) Regulation of proteasome activity in health and disease. Biochim Biophys Acta.

19. Tomko RJ Jr, Hochstrasser M (2013) Molecular Architecture and Assembly of the Eukaryotic Proteasome. Annu Rev Biochem.

20. Bhattacharyya S, Yu H, Mim C, Matouschek A (2014) Regulated protein turnover: snapshots of the proteasome in action. Nat Rev Mol Bio 15: 122–133.

21. Lee B-H, Lee MJ, Park S, Oh D-C, Elsasser S, Chen P-C, Gartner C, Dimova N, Hanna J, Gygi SP, et al. (2010) Enhancement of proteasome activity by a small-molecule inhibitor of USP14. Nature 467: 179–184.

22. Rott R, Szargel R, Haskin J, Bandopadhyay R, Lees AJ, Shani V, Engelender S (2011) α-Synuclein fate is determined by USP9X-regulated monoubiquitination. Proceedings of the National Academy of Sciences 108: 18666–18671.

23. Myeku N, Clelland CL, Emrani S, Kukushkin NV, Yu WH, Goldberg AL, Duff KE (2015) Tau-driven 26S proteasome impairment and cognitive dysfunction can be prevented early in disease by activating cAMP-PKA signaling. Nat Med.

24. Iljina M, Tosatto L, Choi ML, Sang JC, Ye Y, Hughes CD, Bryant CE, Gandhi S, Klenerman D (2016) Arachidonic acid mediates the formation of abundant alpha-helical multimers of alpha-synuclein. Sci Rep 6: 33928.

25. Kristiansen M, Deriziotis P, Dimcheff DE, Jackson GS, Ovaa H, Naumann H, Clarke AR, van Leeuwen FWB, Menéndez-Benito V, Dantuma NP, et al. (2007) Disease-Associated Prion Protein Oligomers Inhibit the 26S Proteasome. Molecular Cell 26: 175–188.

26. Tseng BP, Green KN, Chan JL, Blurton-Jones M, LaFerla FM (2008) Abeta inhibits the proteasome and enhances amyloid and tau accumulation. Neurobiol Aging 29: 1607–1618.

27. Zhang N-Y, Tang Z, Liu C-W (2008) alpha-synuclein protofibrils inhibit 26 S proteasome-mediated protein degradation - Understanding the cytotoxicity of protein protofibrils in neurodegenerative disease pathogenesis. Journal of Biological Chemistry 283: 20288–20298.

28. Wang X, Huang L (2008) Identifying dynamic interactors of protein complexes by quantitative mass spectrometry. Mol Cell Proteomics 7: 46–57.

29. Allen B, Ingram E, Takao M, Smith MJ, Jakes R, Virdee K, Yoshida H, Holzer M, Craxton M, Emson PC, et al. (2002) Abundant tau filaments and nonapoptotic neurodegeneration in transgenic mice expressing human P301S tau protein. Journal of Neuroscience 22: 9340–9351.

30. Kundel F, De S, Flagmeier P, Horrocks MH, Kjaergaard M, Shammas SL, Jackson SE, Dobson CM, Klenerman D (2018) Hsp70 Inhibits the Nucleation and Elongation of Tau and Sequesters Tau Aggregates with High Affinity. ACS Chem Biol 13: 636–646.

31. Brelstaff J, Ossola B, Neher JJ, Klingstedt T, Nilsson KPR, Goedert M, Spillantini MG, Tolkovsky AM (2015) The fluorescent pentameric oligothiophene pFTAA identifies filamentous tau in live neurons cultured from adult P301S tau mice. Front Neurosci 9: 184.

32. Evans LD, Wassmer T, Fraser G, Smith J, Perkinton M, Billinton A, Livesey FJ (2018) Extracellular Monomeric and Aggregated Tau Efficiently Enter Human Neurons through Overlapping but Distinct Pathways. Cell Reports 22: 3612–3624.

33. Ait-Bouziad N, Lv G, Mahul-Mellier A-L, Xiao S, Zorludemir G, Eliezer D, Walz T, Lashuel HA (2017) Discovery and characterization of stable and toxic Tau/phospholipid oligomeric complexes. Nat Commun 8: 1678.

34. Takahashi M, Miyata H, Kametani F, Nonaka T, Akiyama H, Hisanaga S-I, Hasegawa M (2015) Extracellular association of APP and tau fibrils induces intracellular aggregate formation of tau. Acta Neuropathol 129: 895–907.

35. Flagmeier P, De S, Wirthensohn DC, Lee SF, Vincke C, Muyldermans S, Knowles TPJ, Gandhi S, Dobson CM, Klenerman D (2017) Ultrasensitive Measurement of Ca(2+) Influx into Lipid Vesicles Induced by Protein Aggregates. Angew Chem Int Ed 56: 7750–7754.

36. Cremades N, Cohen SIA, Deas E, Abramov AY, Chen AY, Orte A, Sandal M, Clarke RW, Dunne P, Aprile FA, et al. (2012) Direct observation of the interconversion of normal and toxic forms of α-synuclein. Cell 149: 1048–1059.

37. Shorter J, Lindquist S (2004) Hsp104 catalyzes formation and elimination of self-replicating Sup35 prion conformers. Science 304: 1793–1797.

38. Nillegoda NB, Kirstein J, Szlachcic A, Berynskyy M, Stank A, Stengel F, Arnsburg K, Gao X, Scior A, Aebersold R, et al. (2015) Crucial HSP70 co-chaperone complex unlocks metazoan protein disaggregation. Nature 524: 247–251.

39. Gao X, Carroni M, Nussbaum-Krammer C, Mogk A, Nillegoda NB, Szlachcic A, Guilbride DL, Saibil HR, Mayer MP, Bukau B (2015) Human Hsp70 Disaggregase Reverses Parkinson’s-Linked α-Synuclein Amyloid Fibrils. Molecular Cell 59: 781–793.

40. Fitzpatrick AWP, Falcon B, He S, Murzin AG, Murshudov G, Garringer HJ, Crowther RA, Ghetti B, Goedert M, Scheres SHW (2017) Cryo-EM structures of tau filaments from Alzheimer’s disease. Nature 547: 185–190.

41. Braun BC, Glickman M, Kraft R, Dahlmann B, Kloetzel P-M, Finley D, Schmidt M (1999) The base of the proteasome regulatory particle exhibits chaperone-like activity : Abstract : Nature Cell Biology. Nat Cell Biol 1: 221–226.

42. Hao R, Nanduri P, Rao Y, Panichelli RS, Ito A, Yoshida M, Yao T-P (2013) Proteasomes activate aggresome disassembly and clearance by producing unanchored ubiquitin chains. Molecular Cell 51: 819–828.

43. Nanduri P, Hao R, Fitzpatrick T, Yao T-P (2015) Chaperone-mediated 26S proteasome remodeling facilitates free K63 ubiquitin chain production and aggresome clearance. J Biol Chem 290: 9455–9464.

44. Guo Q, Lehmer C, Martínez-Sánchez A, Rudack T, Beck F, Hartmann H, Pérez-Berlanga M, Frottin F, Hipp MS, Hartl FU, et al. (2018) In Situ Structure of Neuronal C9orf72 Poly-GA Aggregates Reveals Proteasome Recruitment. Cell 172: 696–705.e12.

45. Thomas SN, Funk KE, Wan Y, Liao Z, Davies P, Kuret J, Yang AJ (2012) Dual modification of Alzheimer’s disease PHF-tau protein by lysine methylation and ubiquitylation: a mass spectrometry approach. Acta Neuropathol 123: 105–117.

46. Morris M, Knudsen GM, Maeda S, Trinidad JC, Ioanoviciu A, Burlingame AL, Mucke L (2015) Tau post-translational modifications in wild-type and human amyloid precursor protein transgenic mice. Nat Neurosci 18: 1183–1189.

47. Hasegawa M, Fujiwara H, Nonaka T, Wakabayashi K, Takahashi H, Lee VMY, Trojanowski JQ, Mann D, Iwatsubo T (2002) Phosphorylated α-Synuclein Is Ubiquitinated in α-Synucleinopathy Lesions. Journal of Biological Chemistry 277: 49071–49076.

48. Tofaris GK, Razzaq A, Ghetti B, Lilley KS, Spillantini MG (2003) Ubiquitination of alpha-synuclein in Lewy bodies is a pathological event not associated with impairment of proteasome function. Journal of Biological Chemistry 278: 44405–44411.

49. Barrett PJ, Timothy Greenamyre J (2015) Post-translational modification of α-synuclein in Parkinson’s disease. Brain Res 1628: 247–253.

